# CRISPR interference in a *Streptococcus agalactiae* Multi-locus Sequence Type 17 Strain

**DOI:** 10.1101/2025.08.27.672580

**Authors:** William D. Cutts, Aidan W. Flanagan, Brice Gorman, Audrey Sweten, Bryan Estrada, Vishwas Subash, Benjamin Klemp, Kathryn Seely, Austin Sandobal, Katelin Stilen, Taksh Vaghela, Areebah Mehvish, Jacob F. Wood, Alexus Govert, Kristin Hobson, Gideon H. Hillebrand, Thomas A. Hooven, Brandon J. Kim

## Abstract

Group B Streptococcus (GBS), a common colonizer of the human genital and gastrointestinal tracts, is a leading cause of neonatal bacterial meningitis, which can lead to severe neurological complications. The hypervirulent serotype III, sequence type 17 (ST-17) strain COH1 is strongly associated with late-onset disease due to its unique set of virulence factors. However, genetic manipulation of ST-17 strains is notoriously challenging, limiting the ability to study key pathogenic genes. In this study, we developed a CRISPR interference (CRISPRi) system utilizing an endogenous catalytically inactivated Cas9 (dCas9) in the COH1 strain, enabling targeted and tunable gene expression knockdown. We confirmed the efficacy of this system through hemolysis assays, qPCR transcriptional analysis, and in vitro infection models using human brain endothelial cells. The CRISPRi system successfully produced phenotypic knockdowns of essential virulence genes, including *pilA, srr2*, and *iagA*, reducing adhesion, invasion, and inflammatory responses at the blood-brain barrier. This platform enables rapid gene knockdowns for functional genomics in ST-17 GBS, enabling high-throughput screening and pathogenesis research.

**Importance:** Group B Streptococcus (GBS) remains the world’s leading cause of neonatal meningitis. GBS-host interactions at the blood-brain barrier (BBB) are dependent on bacterial factors, including surface factors and two-component systems. Multi-locus sequence type 17 (ST-17) GBS strains are highly associated with neonatal meningitis, and these strains harbor many virulence factors for infection at the BBB. Historically, these factors have been studied using traditional knockout mutagenesis, which has proven challenging in the most common ST-17 lab strain, COH1. This study utilizes CRISPR interference (CRISPRi) to generate rapid expression knockdown. This study validates a CRISPRi-enabled COH1 dCas9 strain as a versatile tool for probing GBS pathogenesis at the BBB.

## Introduction

Group B Streptococcus (GBS), also known as *Streptococcus agalactiae*, is a Gram-positive bacterium that asymptomatically colonizes the genital and gastrointestinal tracts of 20-30% of healthy individuals (1, 2). However, during or shortly after birth, GBS can opportunistically infect neonates and infants, leading to conditions such as sepsis, pneumonia, or meningitis (3–6). Worldwide, GBS is the leading cause of neonatal bacterial meningitis, which is uniformly fatal without medical intervention (1, 2). While modern medical interventions have transformed GBS meningitis from a uniformly fatal illness to an often curable one, mortality remains at 5% to 10%, with survivors facing long-term neurological sequelae such as blindness, deafness, seizure, and stroke (1, 7–9). Additionally, GBS can asymptomatically spread to other body sites and cause disease in adults, including soft tissue infections, bacteremia, urinary tract infections, endocarditis, and meningitis (3–5). Neonatal invasive GBS disease is classified into two types: early-onset disease (EOD), which occurs within the first seven days of life, and late-onset disease (LOD), which develops between 7 and 90 days of life. 97% of neonatal invasive GBS diseases are caused by serotypes I–V, and serotype III accounts for 43% of early-onset disease and 73% of late-onset disease (1, 5, 8). LOD presentation is most commonly meningitis, but can also present as urinary tract, joint, bone, and soft tissue infection, as well as pneumonia and bacteremia (8, 10, 11).

Some GBS strains are more strongly associated with neonatal disease than others. Many strains that fall within serotype III sequence type 17 (ST-17), considered a hypervirulent sequence type, are significantly associated with neonatal meningitis and particularly LOD (5, 8). A significant contributing factor to ST-17’s hypervirulence is the number of adhesins and virulence factors shared across this clade. These include the serine-rich repeat proteins (Srr-1/2), GBS pilus tip adhesin (PilA), hypervirulent GBS adhesin (HvgA), laminin binding protein (Lmb), streptococcal fibrinogen binding protein A (SfbA), and Group B streptococcal surface protein C (BspC), as well as the two-component systems CovR/S and CiaR/S, which have all been described previously through traditional allelic-exchange mutagenesis or transposon mutagenesis (2, 12–24). However, mutagenesis can prove laborious and time-consuming, particularly in ST-17 strains such as COH1, which are notoriously difficult to manipulate.

GBS utilizes an endogenous Type II-A CRISPR-Cas9 system, much like *S. pyogenes*, and this system has recently been used to screen candidate genes using CRISPR interference (CRISPRi) (25). This process involves site-directed mutagenesis of key catalytic residues RuvC-like and HNH domains (D10 & H845, respectively), which yields the ability to produce targeted, tunable expression knockdown in any given candidate gene containing the appropriate NGG protospacer adjacent motif (PAM) sequence (25, 26). Until now, this has only been done in serotype Ia and serotype V strains, which, while valid for their backgrounds, are not as translatable to study at the blood-brain barrier (BBB). We hypothesize that such a system in COH1 will provide a faster path to *in vitro* loss-of-function phenotypes, facilitating the speed of research on genes of interest. We report here the use of our hCMEC/D3 brain endothelial cell model to study infection, specifically quantification of bacterial adhesion, invasion, and chemokine expression after knocking down various virulence genes. Additionally, in the process of generating the COH1 dCas9 mutant for this paper, we report the first use of a Cas12a-based system in the generation of a markerless point mutation in GBS. These findings will be of use in facilitating GBS research.

## Results

### Generation of *Streptococcus agalactiae* str. COH1 dCas9

To utilize CRISPRi in GBS, we opted to make use of the endogenous GBS *cas9* gene, point-mutating it to generate a COH1 strain with catalytically deadened Cas9 (dCas), into which we transform a single plasmid expressing the single guide RNA (sgRNA) used to target dCas9 to the site of interest in the genome (Figure 2A) (25, 27). The mutant was generated using two approaches, with the first point mutation inactivating the HNH-like domain (H845A) generated using classical allelic exchange methods, and the second point mutation inactivating the RuvC-like domain (D10A) generated with a Cas12a-based gene deletion method (25, 26, 28, 29). The Cas12a mutagenesis process required targeting of the genome for cutting, and presentation of a repair cassette on pGBSedit that carried our target (D10A) mutation as well as silent mutations altering the sgRNA target site and PAM to protect any successfully edited mutants (**Figure 1A**). For these silent mutations, we selected codons frequently used by GBS to avoid introducing rare codons that might alter gene expression (30). This is the first instance of this system being used to generate a markerless point mutation in GBS (**Figure 1A**). Mutagenesis was confirmed via Oxford Nanopore whole-genome sequencing (Plasmidsaurus) to verify the correct genotype (**Figure 1B**) (31). No growth kinetics changes were observed with this mutant (**Supp. Figure 1**).

**Figure 1:**
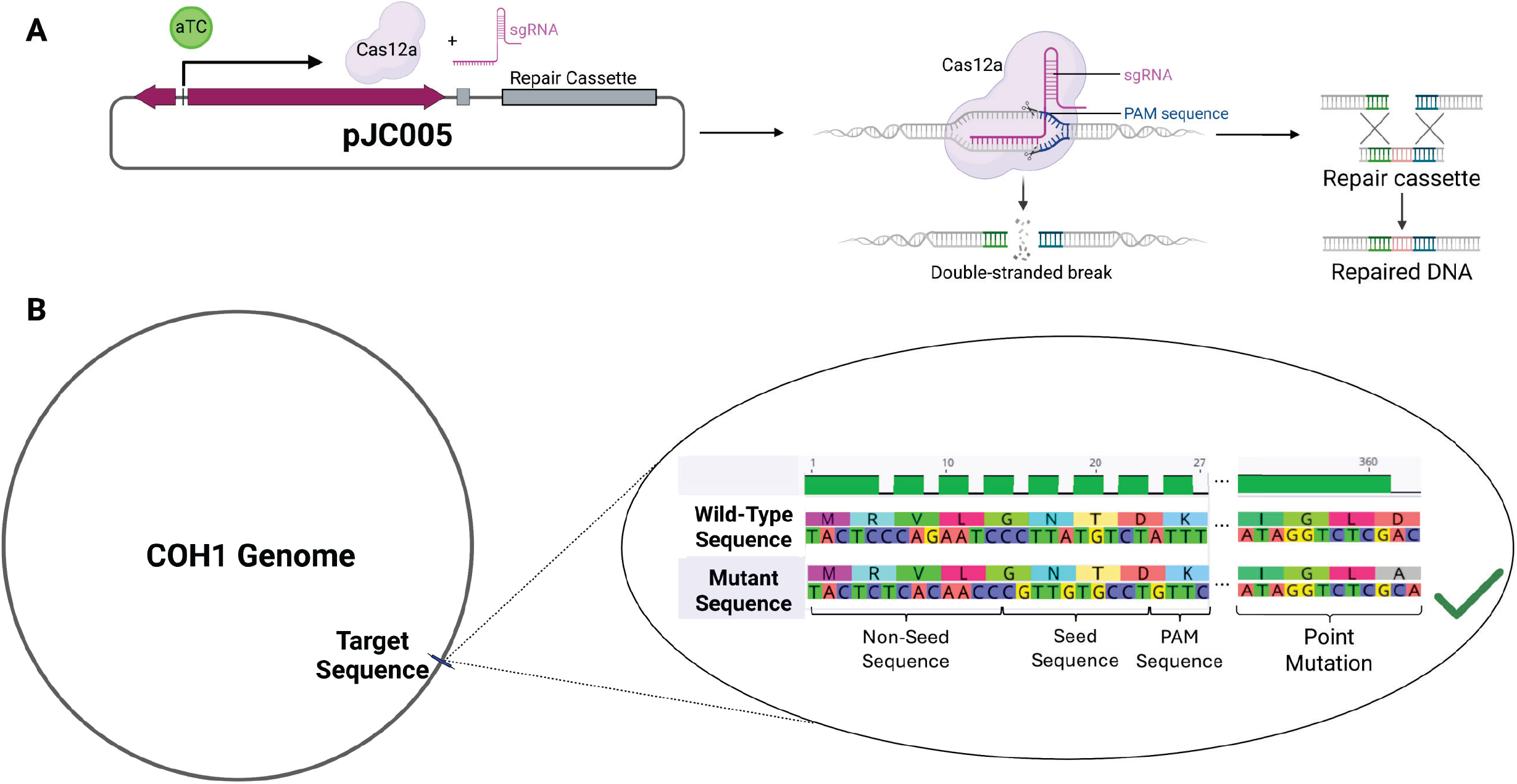
Cas12a-based Point Mutation for CRISPRi. (A) Graphical representation of GBS point mutation process, showing anhydrotetracycline induction of Cas12a and sgRNA expression on the pGBSedit (pGBSedit) plasmid. Also pictured is the inclusion of a repair cassette, containing the homology needed for repair and carrying the desired mutation. (B) Depiction of the genotypic changes needed to generate a point mutation, showing both a hypothetical point mutation (D360A) as well as the silent mutations in the sgRNA target site necessary to protect the plasmid and mutant strain from Cas12a cutting. These silent mutations must be designed on the plasmid repair cassette for the mutation process to be successful.

**Figure 2:**
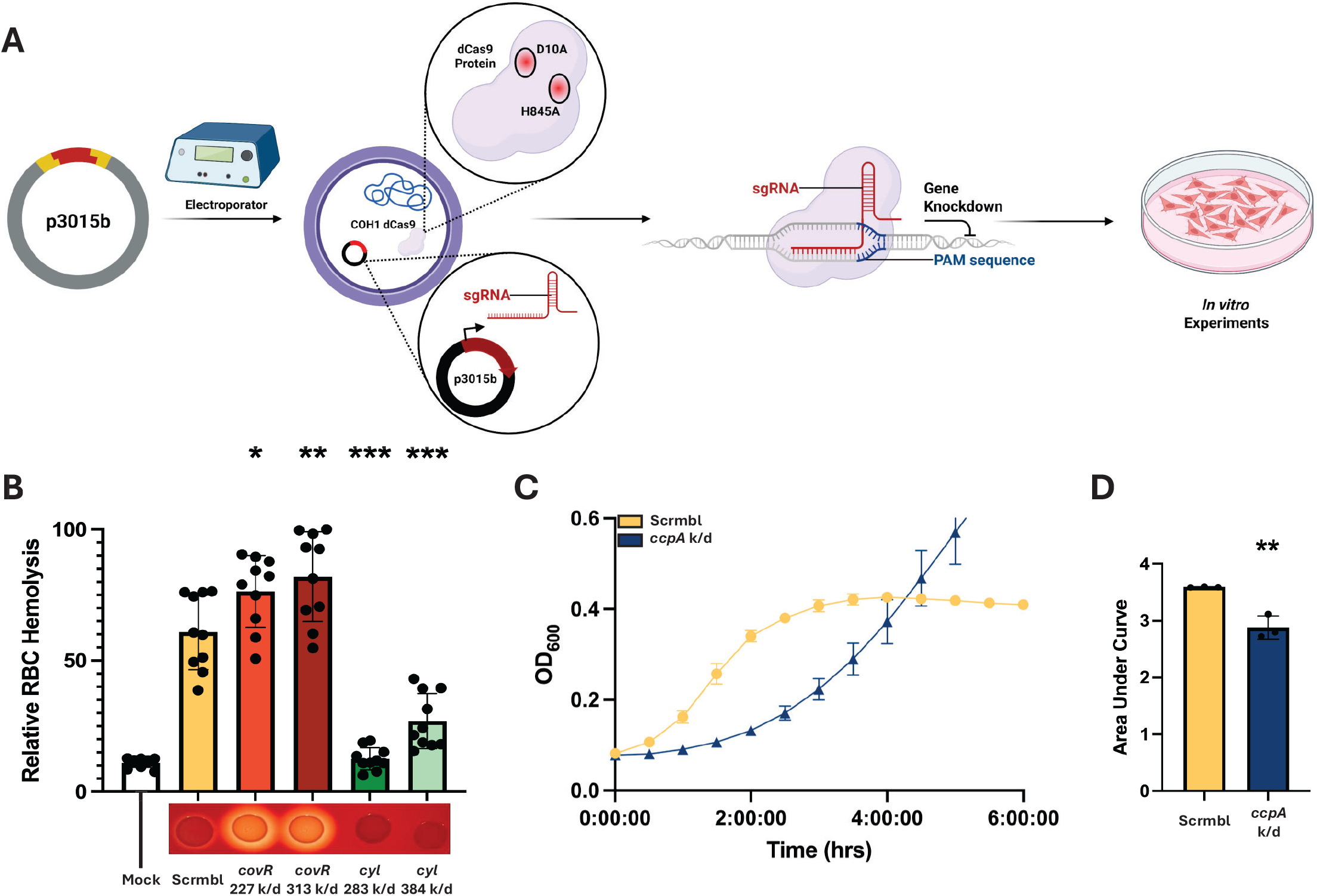
Verification of Phenotype Changes. (A) Graphical depiction of the knockdown generation process. sgRNAs are cloned into the shuttle vector p3015b, then electroporated into *Streptococcus agalactiae* str. COH1 dCas9. These transformants should now have a knockdown phenotype and can be used for subsequent *in vitro* experimentation. (B) Hemolysis assay data, with Mock being PBS-treated sheep’s blood, Scramble being a control infection lacking a targeted sgRNA, as well as two variants of knockdowns of *covR* and the *cyl* operon. Beneath each bar graph is a visual representation of the hemolytic phenotype of each transformant using bacterial growth on sheep’s blood agar, with the *covR* knockdowns showing more hemolysis relative to the control and the *cyl* knockdowns showing less hemolysis. One-way ANOVA was performed, **P < 0.01 and ***P < 0.001. All experiments were performed in biological and technical triplicate, (n=9).

### Verification of Knockdown Phenotypes

To confirm that the mutant strain would show an altered phenotype when targeted with an sgRNA, we performed a hemolysis assay utilizing sgRNAs targeted to the *cyl* operon and the known virulence repressor *covR*, which regulates *cyl* expression (2). To verify reduced hemolytic activity, a 1% red blood cell (RBC) solution was aliquoted into a 96-well plate, infected, and then combined with a PBS suspension of a GBS + sgRNA transformant. Two guides were used for each target gene to evaluate the potential for modulation of the knockdown phenotype. Control groups were instead treated with an equal volume of PBS or with.1% TritonX-100, as well as an experimental control carrying a “scramble” sgRNA sequence that lacks a genomic target in COH1 and should therefore produce no change in phenotype. Triton-X is a detergent that will lyse the RBCs nearly completely, providing a maximum lysis level to normalize data. After a one-hour incubation, cell debris was pelleted, and supernatant was transferred to a new 96-well plate, and absorbance was read at 415 nm to read hemoglobin release as a proxy for the degree of RBC lysis. Results were normalized to the reads of Triton-X-treated samples. Almost all the GBS-infected wells showed some degree of lysis compared to PBS mock wells. The strains harboring *cyl*-targeted guides showed reduced hemolysis relative to the scramble control, and the *covR*-targeted strains showed increased lysis, demonstrating that the guides and CRISPRi system produce expected phenotypes (**Figure 2B**).

Next, to verify the possibility of generating knockdowns of essential genes, a noted strength of CRISPRi, an sgRNA was produced to target the essential gene *ccpA*. After the transformation of this sgRNA into GBS, a growth curve was performed to verify a change in growth kinetics, and the difference was further quantified using an area under the curve (AUC) analysis. Again, we achieved the expected phenotype, as the bacterial lag phase was extended by multiple hours (**Figure 2C**), and area under the curve analysis showed a reduction in AUC (**Figure 2D**). This further validates the CRISPRi system for reducing essential gene function without being lethal to the bacteria (25, 26).

Taken together, the CRISPRi system produced targeted loss-of-function phenotypes as intended.

### qPCR Verification of Transcriptional Alteration

As CRISPRi reduces expression at the transcript level, qPCR was used to verify that the observed phenotypes were attributed to a reduction in targeted mRNA abundance. RNA was isolated via bead beating and standard lysis protocols, and cDNA was synthesized to be used in a qPCR reaction. Knockdown verification was performed using the previously mentioned genes *covR, cylE*, and *ccpA*, as well as in the established virulence factors *srr2, iagA*, and *pilA*. In each case, sgRNA targeted genes showed reduced expression levels (**Figure 3**), though some variability was apparent, with distance from transcriptional start not always trending with expression reduction. The primers used are reported in Table S1.

**Figure 3:**
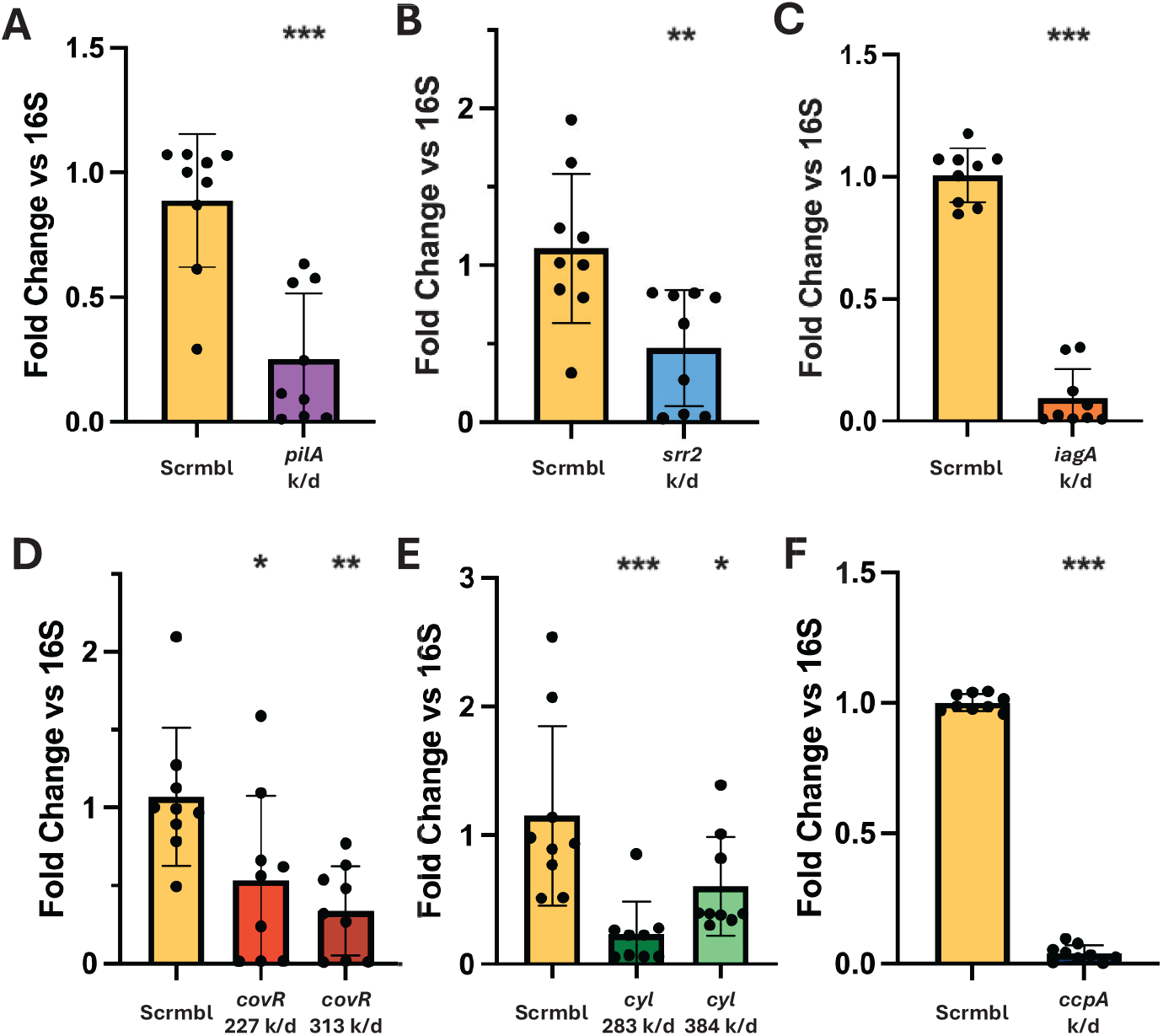
Transcriptional Changes. (A-F) qPCR analysis of target genes following sgRNA transformation into *Streptococcus agalactiae* str. COH1 dCas9 for conformation of CRISPR interference expression modulation. sgRNAs shown include the established virulence genes *pilA* (A), *srr2* (B), *iagA* (C), *covR* (D), and *cyl* (E), as well as the essential gene *ccpA* (F). (A-F). One-way ANOVA was performed, *P < 0.05, **P < 0.01, and ***P < 0.001. All experiments were performed in biological and technical triplicate, (n=9).

### CRISPRi for *in vitro* Infection Studies

As GBS COH1 is associated with neonatal meningitis, we sought to ensure that knockdowns of known factors in the CRISPRi strain would produce reproducible infection phenotypes in established *in vitro* blood-brain barrier modeling systems. For adhesion/invasion assays and cell immune response verification, the hCMEC/D3 cell line was used. For investigating the efficacy of the CRISPRi model in infection, sgRNAs were designed to target the known GBS virulence factors *pilA*, s*rr2*, and i*agA*. These genes all have verified roles in infection at the BBB *in vitro* and with *in vivo* mouse models, as well as further investigation of the molecular mechanisms of these roles in infection (2, 14, 23, 24, 32). Adhesion (reported as cell-association) and invasion results were quantified by infecting hCMEC/D3 cells seeded on a 12-well plate for 30 minutes and 4 hours, respectively. All knockdowns exhibited a reduction in both adhesion (**Figure 4A**) and invasion (**Figure 4B**) relative to scramble. To investigate the known contribution of PilA to immune activation, we performed an experiment using mock-infected cells, infected with scramble, or infected with a PilA-targeted knockdown and collected RNA to quantify changes in chemokine and cytokine expression. We found that the scramble control significantly increased chemokine and cytokine expression when compared to mock, and that knocking down PilA reduced the overall induction of these proinflammatory genes (**Figure 4C-E**). Taken together, these results demonstrate that use of the CRISPRi system mimics phenotypes previously described using classical mutagenesis, thereby providing an opportunity to use CRISPRi to identify potential novel targets in the future.

**Figure 4:**
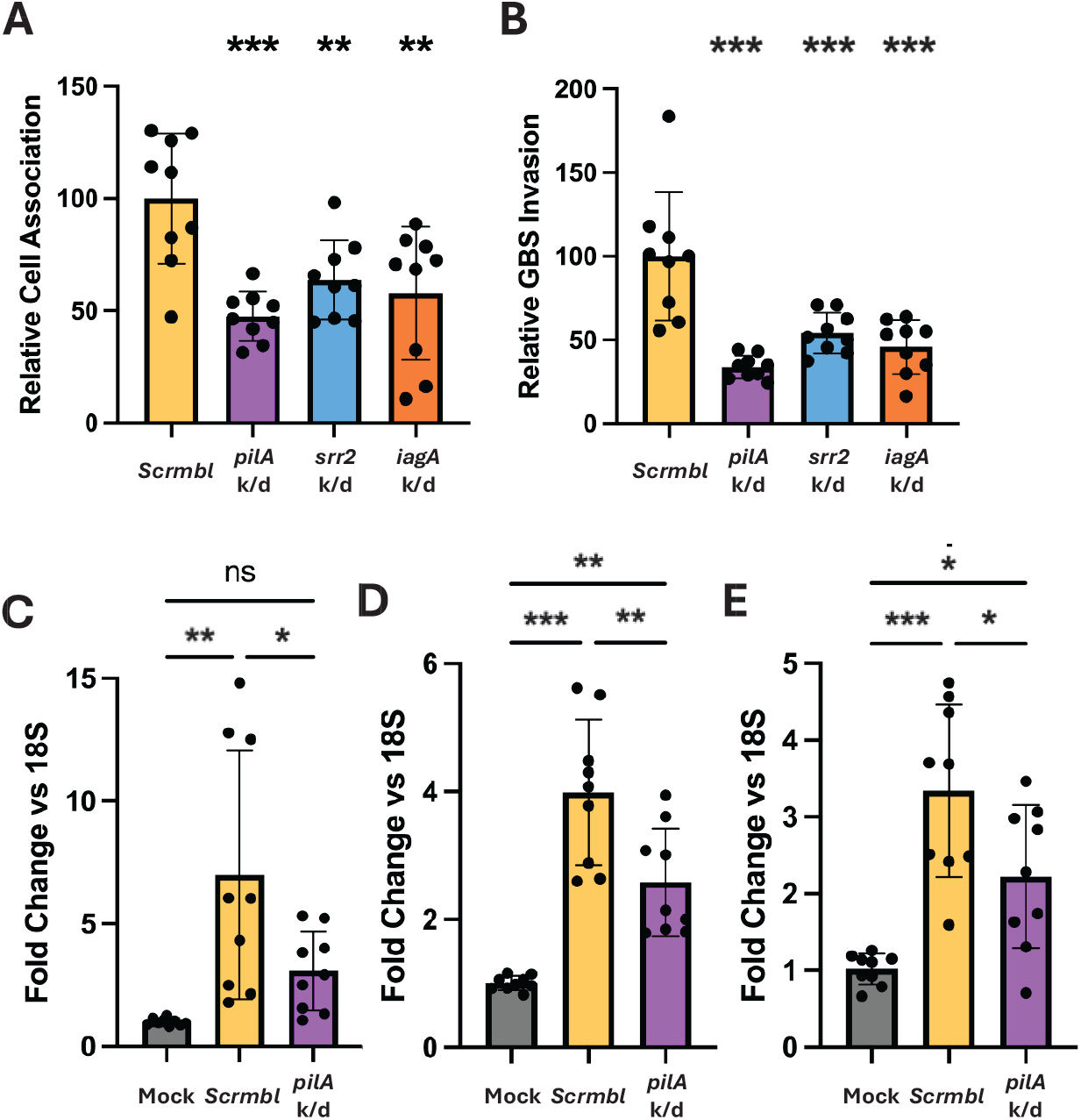
*In vitro* Infection Application. GBS adhesion (A) and invasion (B) rates to hCMEC/D3 monolayers when various well-established virulence genes are knocked down. (C-E) hCMEC/D3 inflammatory marker expression levels when facing challenge with either mock infection, Scramble control, or *pilA* knockdown as a representative virulence gene. Left to right, the qPCR targets are (C) *CXCL8* (IL-8), (D) *CXCL1*, and (E) *CXCL2*. (A-E) One-way ANOVA was performed, *P < 0.05, **P < 0.01, and ***P < 0.001. All experiments were performed in biological and technical triplicate, (n=9).

### *In silico* GBS CRISPRi sgRNA Library Generation

Following the generation of the CRISPRi strain and verification of its utility, we compiled an *in silico* high coverage sgRNA library for use in GBS infection research using CRISPRi. We have designed a double-coverage library of potential sgRNAs using the software tool CHOPCHOP, covering 1944/2073 GBS genes (omitting rRNA and tRNA genes), totaling 3595 sgRNA sequences (Supp. Table 2) to be used as a resource for GBS research. Each of these guides has been generated to minimize off-target effects and consistency of knockdown effects, and evaluated for homology against several other common GBS research strains, including A909, CNCTC 10/84, NEM316, BM110, and CJB111 (Supp. Table 2). This *in silico* library can facilitate future GBS phenotypic research by speeding design of synthetic protospacer oligonucleotides and could potentially inform development of a full-genome coverage knockdown library in GBS.

## Discussion

CRISPRi is an increasingly popular technology for precise, tunable gene regulation, and it has been utilized across a wide range of studies in both eukaryotic and prokaryotic systems. By using a catalytically inactive Cas9 (dCas9) targeted to specific genomic sequences via single-guide RNAs, CRISPRi enables researchers to inhibit transcription without permanently altering the genome, making it a powerful tool for investigating gene function (25, 26, 29, 33).

CRISPRi applications in GBS are advancing, with new tools for GBS rapidly being developed (25, 29). These innovations create opportunities to advance the study of clinically relevant strains such as COH1 as well as the study of complex host-pathogen interactions.

Our main priority in designing this mutant-based endogenous CRISPRi system was ease of use, leveraging tools established in other GBS strains while balancing the knockdown efficiency. To that point, every guide tested in this study produced a viable knockdown phenotype. Notably, these knockdowns were typically achieved, depending on transformation success, within 1-2 weeks from design to performing assays. There was, however, some variability in some of the knockdown phenotypes that is worth noting, particularly in the case of the *covR* knockdowns.

Generally, the results were as expected, with *cyl* operon knockdowns showing reduced or nearly eliminated lytic activity and *covR* knockdowns showing increased lysis relative to the “scramble” control sgRNA group (25, 34–36). Additionally, variation in efficacy from guides was seen. This is expected and desired for regulation purposes via varying the target site distance from the transcriptional start site (TSS), but other variations were observed. For instance, the *cyl* guide targeted 283 bp from TSS showed a greater degree of knockdown than the guide targeted 384 bp from TSS (Figure 2A), in agreement with an established correlation (26). The *covR* guide pair seems to disagree with this; however, more factors may be in play as the less effective *covR* 227 guide targets an AGG PAM sequence, which has been reported elsewhere as being a less efficient PAM sequence in GBS (25).

Also of great importance is the viability of the dCas9 mutant as a tool for the study of COH1 infection modeling *in vitro*. We and others utilize several models to this end, with the iBEC and hCMEC/D3 models increasing the robustness of the findings (37–39). Three well-established virulence genes were selected for knockdown: *pilA, iagA*, and *srr2*, as there is a great deal of literature to which we can compare our knockdown data for verification purposes. The data presented here compares favorably to knockout phenotypes, though less stark in some cases (Figure 4A, B, F) (2, 14, 23, 32). Additionally, we performed qPCR and found that CMEC/D3 cells had a reduced inflammatory response to knockdown strains of virulence factors, with neutrophil recruitment markers (*IL-8, CXCL1*, and *CXCL2)* expressed at lower levels after infection with the pilA knockdown strain compared to the control (Figure 4C–E).

While the system shows strong potential utility, some limitations are worth noting, particularly given the use of a mutated endogenous Cas9 gene rather than purely plasmid-based systems of CRISPRi. The foremost of these is that the system is not placed under an inducible promoter, leaving any knockdown ubiquitously expressed. This limits utility and versatility, as inducible systems would permit a more nuanced experimental design. Additionally, antibiotic selection is necessary to retain the plasmid, which is a limitation for *in vivo* applications. It is also possible that there are unexpected non-transcriptional effects from mutating endogenous *Cas9* rather than a plasmid system. With that said, this is unlikely as previous literature has shown minimal transcriptional change in GBS dCas9 strains relative to wild-type, as PAM scanning is unaffected by this mutation (25). The generation of the *in silico* sgRNA library may be of use to other researchers studying GBS gene function, facilitating faster screening of genes and providing key validation before investing the time and resources to generate mutants. Future innovation on these systems may permit more complex studies, such as CRISPRi-seq, permitting the discovery and interrogation of new genes of interest in GBS COH1.

## Materials and Methods

### Maintenance and differentiation of cell lines

hCMEC/D3 cells were cultured as described previously (40). Cells were maintained on tissue culture flasks coated with 1% rat tail collagen in EndoGRO MV medium (Millipore-Sigma). Cells were grown until 85% confluent, then split for infection experiments. Cells were seeded onto 12-well tissue culture plastic dishes coated with 1% rat tail at 1 × 10^5^ cells/cm^2^ for experiments and allowed to grow to confluency (4 days) at 37°C + 5% CO_2_ before infection experiments were conducted.

### Bacterial strains and growth conditions

Group B Streptococcus (GBS; *Streptococcus agalactiae*) strain COH1 (serotype III, multi-locus sequence type 17 (MLST-17)) was used for this study and cultured in Todd-Hewitt broth (THB) at 37°C in static culture (41, 42). *E. coli* strain DH5α competent cells were used as a reservoir for plasmids, and were grown in Luria-Bertani (LB) broth (LB Miller formulation) at 37°C.

### sgRNA design and protospacer cloning

Guides are designed as described previously to maximize efficiency (26, 33). Great care was taken during design to avoid off-target activity, avoiding seed-sequence homology wherever possible. The software tool CHOPCHOP was used for sgRNA design with that in mind, and NCBI BLAST was used extensively to verify sgRNA efficacy. Guides for *cyl* and *covR* were received from Dr. Thomas Hooven’s lab at the University of Pittsburgh Medical Center, as was the sgRNA shuttle vector p3015b (25). sgRNA sequences are ordered from Eurofins as individual oligos with desired overhangs matching to vector p3015b insertion site, then end phosphorylated (New England Biolabs T4 polynucleotide kinase). Phosphorylated oligos are annealed by heating to 95 °C for 5 minutes, followed by gradual cooling to 4°C to produce the desired spacer. p3015b is miniprepped from DH5α *E. coli* before digestion with BsaI (Eco31I) (Thermofisher). The digested plasmid sample is cleaned (Promega Wizard SV Gel and PCR Cleanup Kit), then the annealed spacer is ligated into the digested shuttle vector (New England Biolabs T4 Ligase) for heat-shock transformation into *E. coli*. Following *E. coli* transformation, the plasmid is miniprepped (ThermoFisher PureLink™ Quick Plasmid Miniprep Kit) for transformation into GBS.

### *In silico* GBS sgRNA Library Design

In general, guides were designed as described above. For library generation, wherever possible, two sgRNAs were generated for every gene (omitting rRNAs and tRNAs) in the COH1 genome. For each gene, one sgRNA was designed near the transcription start site (TSS) (−100 bp to 300 bp from TSS) and another further downstream (>500bp downstream of TSS). Wherever possible AGG PAM sequences were avoided due to the noted reduced efficiency of this PAM in GBS (25). The *silico* tool CHOPCHOP was used for sgRNA generation, as it is designed to help minimize off-target effects. Following sgRNA sequence generation, the best sequences were then checked for homology to other common GBS lab strains (A909, CNCTC 10/84, NEM316, BM110, CJB111), and sgRNAs that would be useful in other strains were chosen wherever possible. This library is provided in Supplemental Table 2.

### GBS competent cell preparation and transformation

Electrocompetent GBS were prepared as described previously, with minor adaptations (43–45). COH1 were grown overnight in a 15 mL THB + 0.6% glycine broth at 37°C. The following day, this culture was diluted in an additional 35 mL of THB + 0.6% Glycine and grown to OD_600_=.6. This culture was pelleted via 4°C centrifugation at 3200 x g and washed twice on ice with a cold 0.6% glycine solution. The remaining pellet was then resuspended in a 400 µL solution of cold 25% PEG 6000 + 10% glycerol and used immediately. Electroporation was also performed as described previously, with only minor alterations (43–45). When electroporating the bacteria, three 3 kV pulses were performed with 5 second intervals between them. Bacteria were then permitted to recover for two hours in THB + 25% PEG 6000 in static culture before plating on THB + 5 µg/mL erythromycin agar plates.

### Generation of GBS dCas9 mutations

Point mutations needed for catalytic deadening of Cas9 were performed in two steps due to the size of the gene and the genomic distance between the two catalytic sites. The H845A missense mutation was generated as previously described using allelic exchange mutagenesis (45). The sucrose-sensitive suicide vector pMBSacB, containing the necessary homology repair cassette with the point mutation, was generously provided by Dr. Thomas Hooven’s lab at the University of Pittsburgh Medical Center. This plasmid was transformed into GBS as described above, and after screening via colony PCR, the transformants were grown overnight in THB + 5ug/mL erythromycin broths at 28 °C. The following day, these broths were passed to fresh tubes of the same broth at 37 °C to begin temperature selection of the first allelic crossover. Following PCR verification of the first allelic crossover event, broths of this strain were passed into broths of THB lacking erythromycin to eliminate selection and permit curing of the plasmid. These broths were repeatedly passed at 28°C and plated onto THB until colony PCR, using primers 750 base pairs up-and-downstream of the homology arm sites, could confirm that a second allelic crossover event occurred. The product of this colony PCR was then inserted into a TOPO TA Vector (TOPO TA Cloning Kit for Sequencing, Thermofisher) and, after being passed in DH5α *E. coli*, submitted to Plasmidsaurus for Oxford Nanopore Sequencing to confirm generation of the first point mutation.

The second point mutation was generated using the Cas12a expressing vector pGBSedit, which was received from Dr. Thomas Hooven’s lab at the University of Pittsburgh Medical Center. A 23 bp protospacer targeted to the Cas9 gene was designed using the appropriate TTTV PAM. This was assembled into pGBSedit (NEB HiFi Assembly 2x Master Mix), and then the homology repair cassette harboring the D10A point mutation was designed and also assembled into the vector. In order to prevent self-cutting of either the plasmid or the final, mutated genome by Cas12, the repair cassette also carries a series of silent mutations in the TTTV PAM used for targeting as well as the targeted sequence itself. The remainder of the process was performed as previously described by Hillebrand et al (29). The plasmid is electroporated into GBS, and, after screening and re-streaking, a colony is inoculated into THB + 5 µg/mL erythromycin broth and grown overnight. This broth is then sub-cultured, and eventually 250ng/mL anhydrotetracycline is added, followed by a one-hour incubation period. The culture is then plated onto THB agar + 5ug/mL erythromycin + 250ng/mL anhydrotetracycline agar plates and allowed to grow overnight. Mutant PCR screening for point mutation is performed by designing a primer to match the silent mutated sgRNA targeting sequence of the mutant gene as the forward primer and using a primer 750 base pairs outside the homology region as the reverse. PCR was performed using the Monserate Midas Quik-Load PCR Mix 2X (2004-G). To ensure specificity for the point mutation, it is necessary to avoid using any polymerase mixes that contain isostabilizing compounds. Raising the annealing temp 3-5°C above the norm for Taq PCR may also be necessary. The PCR-confirmed mutant was then re-streaked, cured of the plasmid by passes in media without erythromycin, and then the genome was sent to Plasmidsaurus for whole-genome sequencing to confirm the desired genotype.

### GBS infection assays

hCMEC/D3 or iBEC cells were seeded onto 12-well plates. Before infection assays, GBS *dcas*9 knockdowns were grown overnight at 37°C in Todd Hewitt Broth (THB), supplemented with 5µg/mL erythromycin to aid in selection for the p3015b vector containing the protospacer. From the overnight culture, bacteria were subcultured into fresh THB + 5µg/mL erythromycin and grown to OD_600_= 0.4-0.6. Bacteria were spun down and resuspended in PBS to OD_600_= 0.4. Bacteria were then diluted 1:10 in EC medium to a multiplicity of infection (MOI) of 10. Adherence and invasion assays were conducted following previously described protocols (46, 47).

For the bacterial adherence assays, bacteria and cells were incubated for 30 minutes at 37^°^C + 5% CO_2_. The cells were then washed 5x with PBS to remove non-adherent bacteria, lysed with 0.025% Triton X-100, diluted in PBS, and plated onto THB + 5µg/mL erythromycin plates. Plates were incubated overnight at 37°C, with colonies being quantified the following day. For the invasion assays, bacteria and cells were incubated for 2 more hours before adding 100 µg/mL gentamicin. After 2 more hours of incubation at 37°C + 5% CO_2_, cells were washed 3x with PBS to remove antibiotics and plated onto THB plates + 5µg/mL erythromycin plates. Plates were incubated overnight as above for quantification. Cell-association and invasion were quantified with the formula 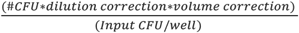 and normalized to MOI association/invasion rates. The true MOI was determined for each experiment.

### Hemolysis assays

Sheep red blood cells (RBCs) were prepared by washing 5 mL of defibrinated blood (Hemostat Labs) with an equal volume of Hank’s Balanced Salt Solution (HBSS) (VWR)at 4°C. The RBCs were pelleted by centrifugation at 500 × g for 15 min at 4°C and resuspended in 5 mL of HBSS. This washing step was repeated three times. A 0.1% Triton X-100 in PBS solution was prepared if not previously made to serve as a positive control. The washed RBCs were then pelleted by centrifugation at 500 × g for 15 min at 4°C. The RBC pellet (50 μL) was diluted into 5 mL HBSS to generate a 1% red blood cell suspension (RBCS).

A 96-well plate was used to set up the assay. Each well contained 100 μL of the 1% RBCS combined with 100 μL of the bacterial PBS suspensions or control solutions. The plate was incubated at 37°C with 5% CO_2_ for 1 hour.

Following incubation, the plate was centrifuged at 2000 rpm for 5 min at 4°C. Supernatants (100 μL) were transferred to new wells on the same 96-well plate, and hemoglobin release was measured by recording absorbance at 415 nm using a spectrophotometer. Percent hemolysis was calculated relative to a positive control consisting of RBCs treated with 0.1% Triton X-100 (100 μL), which represented 100% hemolysis.

### RNA isolation and qPCR

When evaluating human cell inflammatory responses to knockdowns, hCMEC/D3 or iBEC cells were seeded onto 12-well plates and infected with GBS at an MOI of 10. Immediately after 5 hours of infection, the total RNA was collected using the Macherey–Nagel NucleoSpin RNA kit (Macherey–Nagel). cDNA was synthesized with the qScript cDNA Synthesis kit (Quantabio), followed by SYBR Green (PowerUp SYBR Green Master Mix, Thermo Fisher) qPCR for each of the targets: IL-8, CXCL1, CXCL2, primers all listed in Table S1. 18S was used as the housekeeping gene for human cell lines.

For bacterial knockdown qPCR verification, bacteria were grown as described for infection experiments to OD_600_=.4 to ensure mid-log growth phase. Bacteria were resuspended in RA3 lysis buffer, then bead-beaten to lyse before being processed as described above. qPCR data were collected on the QuantStudio3 system (Applied Biosystems), and data are presented as fold change using the delta–delta–CT calculation (48).

## Acknowledgments

B.J.K. is supported by the National Institute of Neurological Disorders and Stroke grant R15NS131921, and the National Institute of Allergy and Infectious Diseases grant R03AI185593 and received salary support from both grants. B.J.K. is also supported by startup funding at the University of Texas at Dallas. T.A.H is supported by National Institutes of Allergy and Infectious Disease (NIAID) R21AI178067, R01AI182835, and R01AI177991. G.H.H. is supported by a UPMC Children’s Hospital of Pittsburgh Research Advisory Council graduate student grant.

T.A.H and G.H.H. contributed the p3015b plasmid and some of the constructs used in this study. W.D.C wrote the manuscript with assistance from J.F.W. and performed all experiments alongside A.W.F, B.G., A.S. and J.F.W. W.D.C., A.W.F., B.G., B.E., V.S., B.K., K.S., K.S., T.V., A.M., A.G., and K.H. all contributed to sgRNA library generation.

## Conflict of Interest Statement

The authors declare that the research was conducted in the absence of any commercial or financial relationships that could be construed as a potential conflict of interest.

## Data availability

The sgRNA library has been made available as supplementary table 2.

